# Cannabidiol prevents spontaneous fear recovery after extinction and ameliorates stress-induced extinction resistance

**DOI:** 10.1101/2022.07.18.500460

**Authors:** Eleni P. Papagianni, William G. Warren, Helen J. Cassaday, Carl W. Stevenson

## Abstract

Cannabidiol, the main non-psychotropic constituent of cannabis, has potential as a treatment for anxiety-related disorders since it reduces learned fear expression and enhances fear extinction. The return of fear over time after successful extinction and stress-induced extinction resistance are potential barriers to the treatment of these disorders with extinction-based psychological therapy. In two experiments using rats subjected to auditory fear conditioning, we determined the effects of systemic cannabidiol treatment on (1) delayed extinction and later spontaneous fear recovery, and (2) extinction resistance caused by immediate extinction (the immediate extinction deficit (IED)). In Experiment 1, cannabidiol was given before delayed extinction occurring 24 hr after conditioning, with extinction recall and spontaneous fear recovery tested drug-free 1 and 21 days after extinction, respectively. We found that cannabidiol had no effect on extinction recall but it prevented spontaneous fear recovery. In Experiment 2, the IED procedure was first validated, with immediate extinction occurring 30 min after conditioning. We confirmed that immediate extinction impaired extinction recall, compared to delayed extinction. Next, cannabidiol was given before immediate or no extinction, with extinction recall tested drug-free the next day. We found that cannabidiol rescued the IED, which did not involve effects on fear memory consolidation. In summary, cannabidiol prevented spontaneous fear recovery after delayed extinction and ameliorated extinction resistance caused by immediate extinction. Although the pharmacological mechanisms underlying these effects remain to be determined, our results add to evidence indicating that cannabidiol might prove useful as an adjunct for potentiating the psychological treatment of anxiety-related disorders.

## Introduction

Various anxiety-related disorders are associated with abnormally persistent fear-related memories involving disturbances in their inhibition through extinction. This type of inhibitory learning reduces learned fear expression and forms the theoretical basis of exposure-based therapy as a psychological treatment for these disorders [1-4]. A promising avenue of preclinical research is investigating the pharmacological potentiation of fear extinction, which models the use of medications as adjuncts for enhancing the efficacy of exposure therapy [5]. Cannabidiol is the main non-psychotropic constituent of cannabis and has anxiolytic potential [6-9]. Acute cannabidiol treatment reduces the expression of learned fear [6, 10-13]. Cannabidiol also enhances fear extinction to facilitate the reduction of learned fear expression [12, 14-16]. This suggests that combining cannabidiol with exposure therapy might be a successful strategy to improve the treatment of anxiety-related disorders.

Although psychological therapies are effective, a considerable proportion of patients do not respond adequately to these treatments. The effects of exposure therapy can also be short-lived, resulting in symptom relapse in the long-term after treatment [5]. Fear can return in various ways after successful extinction, such as with the passage of time through a process known as spontaneous fear recovery [17]. Interestingly, drug treatments that enhance extinction can also protect against the later spontaneous recovery of fear [18-19]. Discovering potential treatments that strengthen extinction to prevent spontaneous fear recovery has translational relevance for mitigating symptom relapse in anxiety-related disorders. Although cannabidiol has been shown to enhance fear extinction, its effect on the later spontaneous recovery of fear remains to be determined.

Investigating the pharmacological facilitation of extinction in preclinical models of extinction resistance is also translationally relevant since aberrant extinction is a feature of anxiety-related disorders [1-4]. Growing evidence indicates that various types of stress impair fear extinction [20-21]. The immediate extinction deficit (IED) refers to the impairment in lasting fear reduction that occurs when extinction is conducted after recent fear conditioning [22]. Previous studies provide evidence that this IED is caused by the high state of arousal or stress resulting from recent fear conditioning. Standard delayed extinction occurring 24 hr after conditioning results in successful extinction recall but presenting footshock immediately before delayed extinction impairs its recall [23]. Moreover, weaker conditioning fails to induce the IED, while the IED that normally occurs with stronger conditioning is rescued via pharmacological blockade of receptor signalling by neuromodulatory and hormonal stress mediators [23-28]. Therefore the IED provides a useful model of extinction resistance that can be used to test potential treatments but the effect of cannabidiol on immediate extinction has not been examined.

In this study we conducted two experiments using rats subjected to auditory fear conditioning to investigate the effects of cannabidiol on the return of fear over time after successful extinction and on stress-induced extinction resistance. In Experiment 1 we examined the effects of cannabidiol given before delayed extinction on extinction recall and later spontaneous fear recovery. In Experiment 2 we first validated the IED procedure and then examined the effects of cannabidiol given before immediate extinction on extinction recall; no extinction controls were also included to determine if any effects of cannabidiol required extinction or involved potential effects on fear memory consolidation.

## Results

### Cannabidiol given before delayed extinction prevents the later spontaneous recovery of fear

The effects of cannabidiol given before delayed extinction on learned fear expression, extinction, and extinction recall and spontaneous fear recovery testing are shown in Fig 1. There were no differences in freezing in response to the tone-shock pairings during fear conditioning between the groups (n=10/group; Fig 1B). Two-way ANOVA revealed no main effect of dose (F _(2, 27)_ = 0.93, P = 0.41) or dose x trial interaction (F _(8, 108)_ = 0.51, P = 0.85). Compared to vehicle, cannabidiol decreased freezing before tone presentations during delayed extinction (Fig 1C). One-way ANOVA revealed a significant main effect of dose (F _(2, 27)_ = 3.42, P = 0.047) and post-hoc analysis showed that freezing was significantly decreased by the 10 mg/kg dose, compared to vehicle (P < 0.05). This indicates that cannabidiol reduced baseline fear expression before delayed extinction. Cannabidiol had no effect on tone-induced freezing at the start of delayed extinction (Fig 1D). One-way ANOVA revealed no main effect of dose (F _(2, 27)_ = 0.98, P = 0.39), indicating a lack of effect of cannabidiol on cued fear expression. Cannabidiol also had no effect on tone-induced freezing during delayed extinction (Fig 1E). Two-way ANOVA revealed no main effect of dose (F _(2, 27)_ = 1.52, P = 0.24) or dose x trial block interaction (F _(18, 243)_ = 1.28, P = 0.20), indicating a lack of effect of cannabidiol on extinction learning. Freezing before tone presentations was increased during spontaneous fear recovery testing, compared to extinction recall testing, but cannabidiol had no lasting effect on freezing before tone presentations during either test (Fig 1F). Two-way ANOVA revealed a significant main effect of time (F _(1, 27)_ = 9.92, P = 0.004) but no main effect of dose (F _(2, 27)_ = 0.87, P = 0.43) or dose x time interaction (F _(2, 27)_ = 1.15, P = 0.33). This indicates that the spontaneous recovery of baseline fear occurred, which was unaffected by cannabidiol. Tone-induced freezing was increased during spontaneous fear recovery testing, compared to extinction recall testing, with vehicle but not with cannabidiol (Fig 1G). Two-way ANOVA revealed a significant main effect of time (F _(1, 27)_ = 4.9, P = 0.036) and dose x time interaction (F _(2, 27)_ = 8.41, P = 0.0014). Post-hoc analysis showed that while cannabidiol had no lasting effect on tone-induced freezing during extinction recall testing, both doses resulted in decreased freezing in response to the tones during spontaneous fear recovery testing, compared to vehicle (P < 0.05). This indicates that cannabidiol prevented the spontaneous recovery of cued fear after delayed extinction.

**Fig 1.**
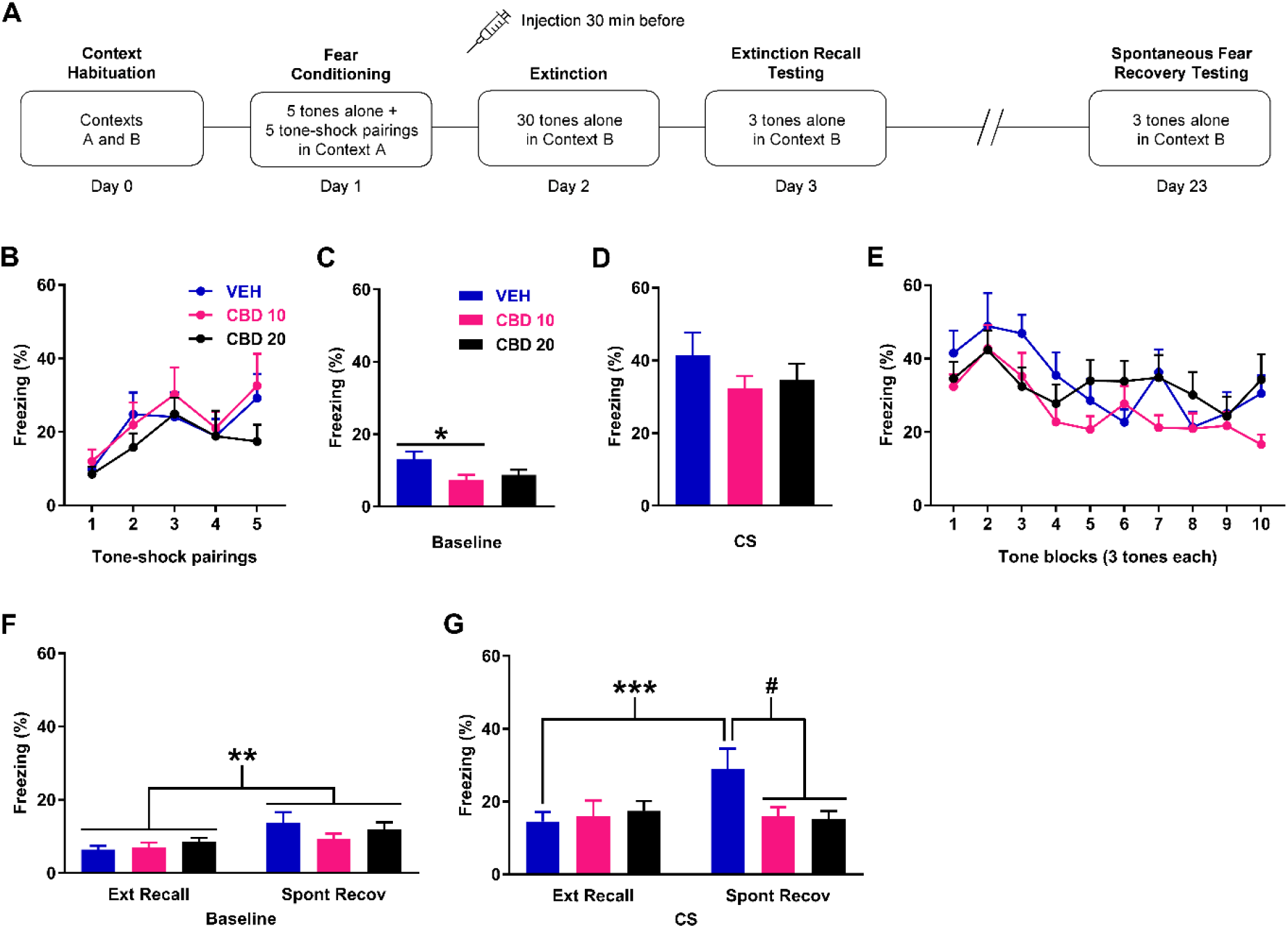
Effects of cannabidiol (CBD) given at 10 mg/kg (CBD 10) or 20 mg/kg (CBD 20) on freezing during delayed extinction, extinction recall testing, and later spontaneous fear recovery testing. A) Schematic representation of the behavioural testing and drug administration procedures used. B) Freezing in response to tone-shock pairings during fear conditioning. There were no differences in freezing between the groups (n=10/group) to receive vehicle (VEH), CBD 10, or CBD 20 before delayed extinction. C) Freezing before tone presentations (Baseline) during delayed extinction. Compared to VEH, CBD 10 decreased freezing (**\*** P < 0.05). D) Freezing in response to the first block of tones at the start of delayed extinction. CBD had no effect on freezing. E) Tone-induced freezing during delayed extinction. CBD had no effect on freezing. F) Freezing before tone presentations (Baseline) during extinction recall (Ext Recall) and spontaneous fear recovery (Spont Recov) testing. Compared to Ext Recall testing, freezing was increased during Spont Recov testing (**\*\*** P < 0.01). CBD resulted in no effect on freezing. G) Tone-induced freezing during Ext Recall and Spont Recov testing. Freezing was increased during Spont Recov testing, compared to Ext Recall testing, with VEH (**\*\*\*** P < 0.001) but not with CBD 10 or CBD 20. There was no effect of CBD on freezing during Ext Recall testing. Compared to VEH, CBD 10 or CBD 20 decreased freezing during Spont Recov testing (**#** P < 0.05).

### Immediate extinction results in impaired extinction recall in comparison to delayed extinction

Validation of the IED procedure, which compared between the effects of immediate and delayed extinction on extinction recall, is shown in Fig 2. There were no differences in freezing in response to the tone-shock pairings during fear conditioning between the groups (n=10/group; Fig 2B). Two-way ANOVA revealed no main effect of group (F _(3, 36)_ = 0.20, P = 0.89) or group x trial interaction (F _(12, 144)_ = 0.45, P = 0.94). There were no differences in freezing before tone presentations during extinction (or no extinction) between the delayed (extinction and no extinction combined) and immediate (extinction and no extinction combined) groups (t _(38)_ = 0.82, P = 0.42), indicating similar baseline fear expression in both groups (Fig 2C). There were also no differences in freezing at the start of extinction between delayed and immediate extinction (t _(18)_ = 0.45, P = 0.67), indicating similar cued fear expression in both groups (Fig 2D). Similarly, there were no differences in tone-induced freezing during extinction between delayed and immediate extinction (Fig 2E). Two-way ANOVA revealed no main effect of group (F _(1, 18)_ = 0.60, P = 0.45) or group x trial block interaction (F _(8, 144)_ = 0.29, P = 0.97), indicating similar extinction learning with delayed and immediate extinction. Freezing before tone presentations during extinction recall testing was increased with immediate extinction, compared to delayed no extinction (Fig 2F). One-way ANOVA revealed a significant main effect of group (F _(3, 36)_ = 3.53, P = 0.024) and post-hoc analysis confirmed this difference to be significant (P < 0.05), indicating that baseline fear expression before extinction recall testing was higher with immediate extinction than with delayed no extinction. Tone-induced freezing during extinction recall, based on mean freezing across the five tones, was increased with immediate extinction in comparison to delayed extinction (Fig 2G). One-way ANOVA revealed a significant main effect of group (F _(3, 36)_ = 3.81, P = 0.018), which post-hoc analysis confirmed to be significant (P < 0.05). This indicates that immediate extinction resulted in impaired extinction recall, compared to delayed extinction. However, post-hoc analysis also showed no differences in freezing between immediate extinction and the no extinction controls (P > 0.05), which led us to examine freezing in response to each tone (Fig 2H). Two-way ANOVA revealed a significant group x trial interaction (F _(12, 144)_ = 14.87, P < 0.0001) and post-hoc analysis showed that freezing was significantly decreased with delayed extinction, compared to immediate extinction and the no extinction controls, during tones 4-5 (P < 0.001). This indicates that extinction recall was impaired with immediate extinction, compared to delayed extinction, later on during testing and confirms the successful validation of the IED procedure.

**Fig 2.**
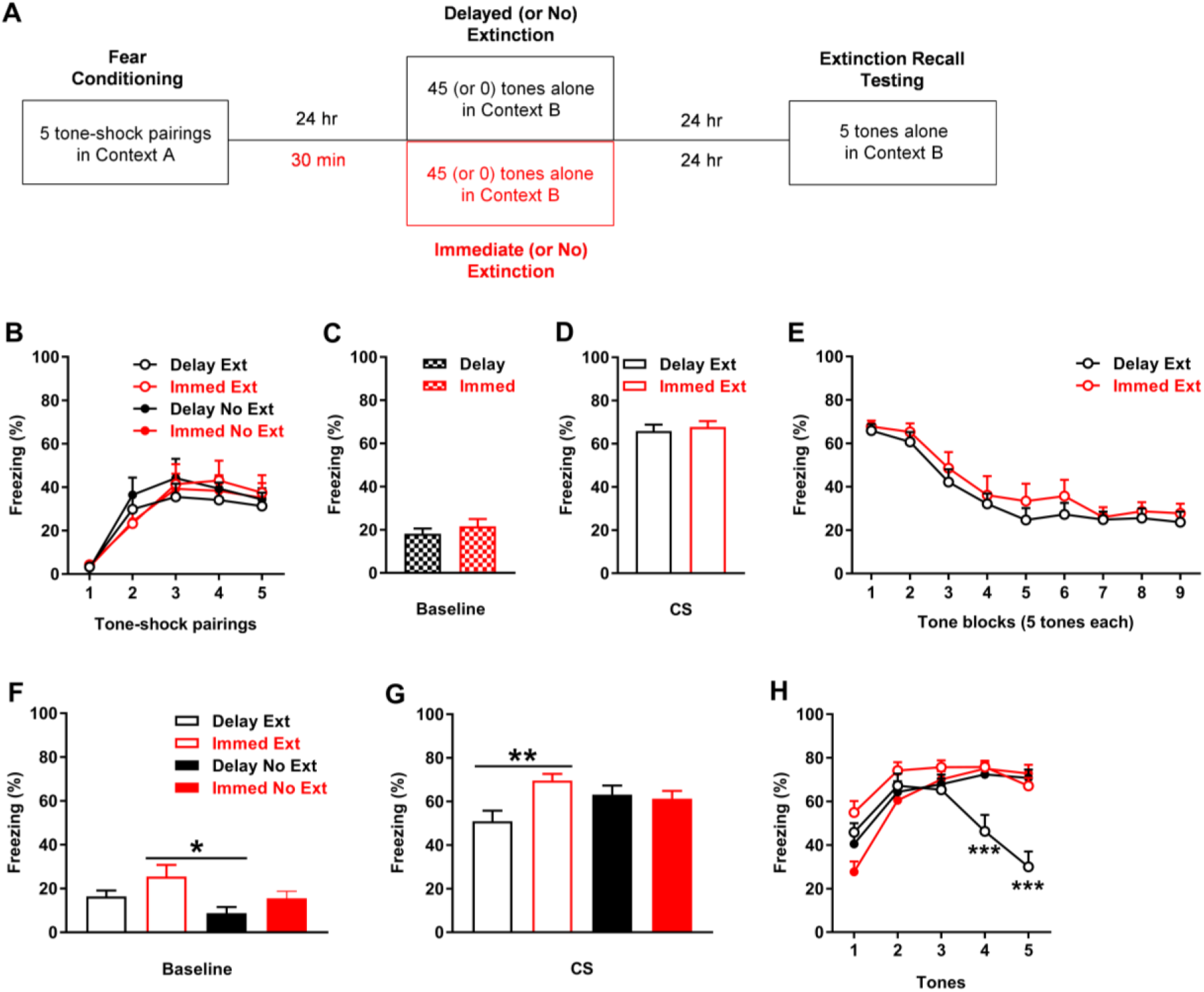
Validation of the immediate extinction deficit procedure. A) Schematic representation of the behavioural testing procedures used. B) Freezing in response to tone-shock pairings during fear conditioning. There were no differences in freezing between the groups (n=10/group) to later undergo delayed (Delay) or immediate (Immed) extinction (Delay Ext, Immed Ext) or no extinction (Delay No Ext, Immed No Ext). C) Freezing before tone presentations (Baseline) during Delay or Immed (or no) extinction. There were no differences in freezing between Delay (Ext and No Ext combined) and Immed (Ext and No Ext combined). D) Freezing in response to the first block of tones at the start of Delay Ext or Immed Ext. There were no differences in freezing between Delay Ext and Immed Ext. E) Tone-induced freezing during Delay Ext and Immed Ext. There were no differences in freezing between Delay Ext and Immed Ext. F) Freezing before tone presentations (Baseline) during extinction recall. Freezing was increased with Immed Ext, compared to Delay No Ext (* P < 0.05). G) Tone-induced freezing across extinction recall testing. Freezing was increased with Immed Ext, compared to Delay Ext (* P < 0.05). H) Freezing in response to each tone during extinction recall testing. Freezing was decreased with Delay Ext, compared to the other groups, during tones 4-5 (*** P < 0.001).

### Cannabidiol given before immediate extinction rescues the immediate extinction deficit

The effects of cannabidiol given before immediate extinction on learned fear expression, immediate extinction, and extinction recall testing are shown in Fig 3. There were no differences in freezing in response to the tone-shock pairings during fear conditioning between the groups (n=10/group; Fig 3B). Two-way ANOVA revealed no main effect of group (F _(3, 36)_ = 0.74, P = 0.54) or group x trial interaction (F _(12, 144)_ = 0.66, P = 0.78). There were no differences in freezing before tone presentations during immediate (or no) extinction between the vehicle (extinction and no extinction combined) and cannabidiol (extinction and no extinction combined) groups (t _(38)_ = 0.16, P = 0.88), indicating a lack of effect of cannabidiol on baseline fear expression (Fig 3C). Compared to vehicle, cannabidiol decreased tone-induced freezing at the start of immediate extinction (t _(18)_ = 3.10, P = 0.0062), indicating that cannabidiol reduced cued fear expression (Fig 3D). Cannabidiol had no effect on tone-induced freezing during immediate extinction (Fig 3E). Two-way ANOVA revealed no main effect of group (F _(1, 18)_ = 3.72, P = 0.07) or group x trial block interaction (F _(8, 144)_ = 0.96, P = 0.47), indicating a lack of effect of cannabidiol on extinction learning. Freezing before tone presentations during extinction recall testing was increased with vehicle extinction, compared to cannabidiol extinction and both no extinction controls (Fig 3F). One-way ANOVA revealed a significant main effect of group (F _(3, 36)_ = 5.12, P = 0.0047) and post-hoc analysis confirmed that these differences were significant (P < 0.05). This indicates that baseline fear expression before extinction recall testing was higher with vehicle extinction, compared to the other groups. Tone-induced freezing during extinction recall, based on mean freezing across the 10 tones, was decreased with cannabidiol extinction in comparison to the other groups (Fig 3G). One-way ANOVA revealed a significant main effect of group (F _(3, 36)_ = 8.03, P = 0.0003) and post-hoc analysis confirmed that these differences were significant (P < 0.01), while also showing no differences between the no extinction controls (P > 0.05). This was confirmed by examining freezing in response to each tone (Fig 3H). Two-way ANOVA revealed a significant group x trial interaction (F _(27, 324)_ = 6.74, P < 0.0001) and post-hoc analysis showed that freezing was significantly decreased with cannabidiol extinction, compared to vehicle extinction and the no extinction controls, during tones 4-7 and 10 (P < 0.05). Freezing was also significantly decreased with vehicle extinction, compared to both no extinction controls, during tones 7-10 (P < 0.05). However, there were no differences in freezing between vehicle no extinction and cannabidiol no extinction throughout extinction recall testing (P > 0.05). This indicates that extinction recall was enhanced by cannabidiol extinction, compared to vehicle extinction, and shows that cannabidiol rescued the IED. It also indicates that this effect of cannabidiol required extinction and rules out the possibility that this involved an effect on fear memory consolidation, given the lack of effect of cannabidiol with no extinction. Finally, the decrease in freezing with vehicle extinction, compared to the no extinction controls, later on during extinction recall testing indicates that savings of extinction occurred with immediate extinction.

**Fig 3.**
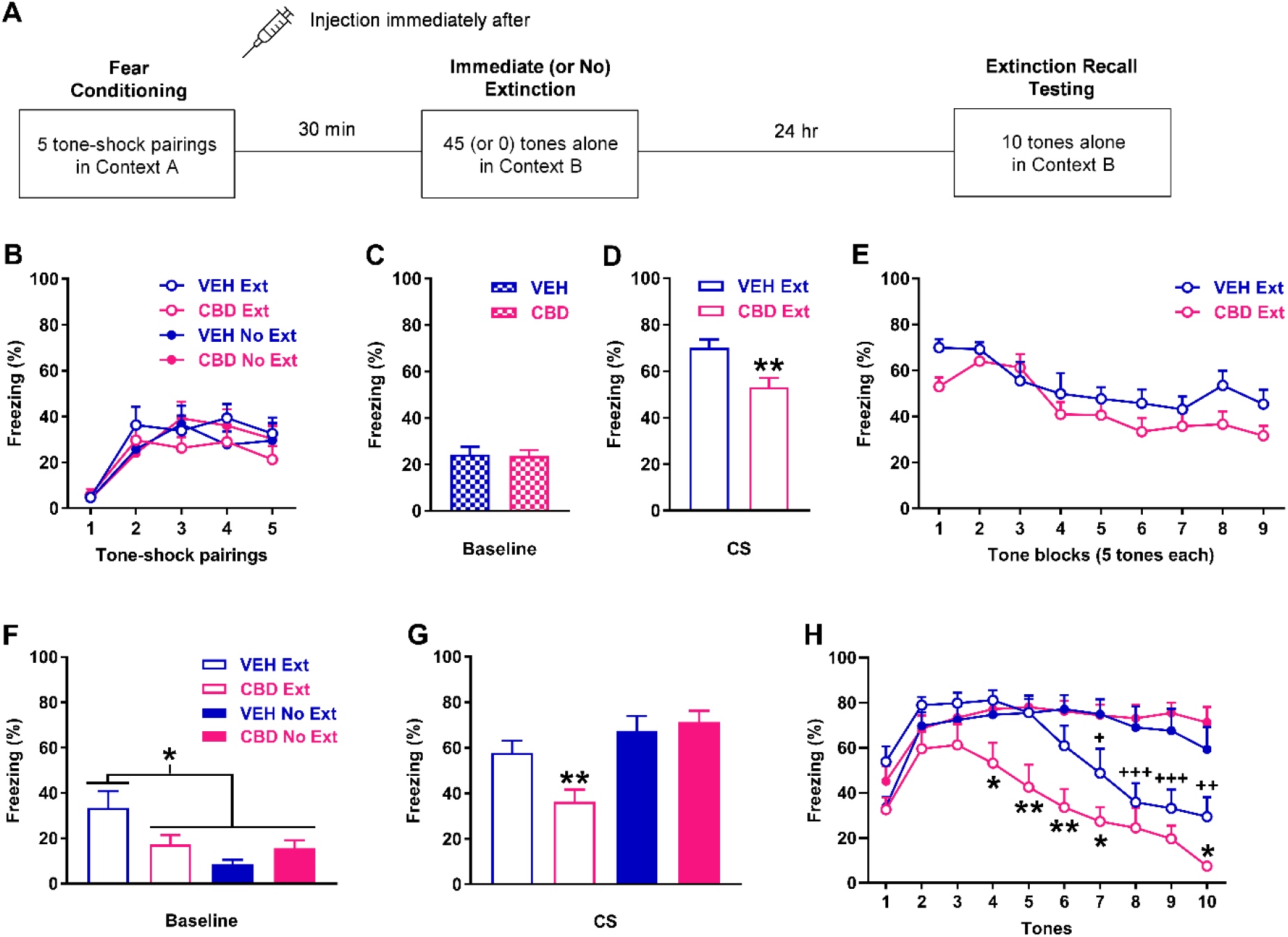
Effects of cannabidiol (CBD) on freezing during immediate extinction and extinction recall testing. A) Schematic representation of the behavioural testing and drug administration procedures used. B) Freezing in response to tone-shock pairings during fear conditioning. There were no differences in freezing between the groups (n=10/group) to receive vehicle (VEH) or CBD before immediate extinction (VEH Ext, CBD Ext) or no extinction (VEH No Ext, CBD No Ext). C) Freezing before tone presentations (Baseline) during immediate (or no) extinction. There were no differences in freezing between VEH (Ext and No Ext combined) and CBD (Ext and No Ext combined). D) Freezing in response to the first block of tones at the start of immediate extinction. Compared to VEH Ext, CBD Ext decreased freezing (\***\*** P < 0.01). E) Tone-induced freezing during immediate extinction. CBD had no effect on freezing. F) Freezing before tone presentations (Baseline) during extinction recall. Freezing was increased with VEH Ext, compared to the other groups (* P < 0.05). G) Tone-induced freezing across extinction recall testing. Freezing was decreased with CBD Ext, compared to the other groups (** P < 0.01). H) Freezing in response to each tone during extinction recall testing. Freezing was decreased with CBD Ext, compared to the other groups, during tones 4-7 and 10 (* P < 0.05, ** P < 0.01). Freezing was also decreased with VEH Ext, compared to VEH No Ext and CBD No Ext, during tones 7-10 (+ P < 0.05, ++ P < 0.01, +++ P < 0.001).

## Discussion

In Experiment 1 we found that systemic cannabidiol treatment before delayed extinction reduced baseline fear expression acutely without affecting cued fear expression or extinction learning. Cannabidiol had no lasting effect on extinction recall but it prevented the later spontaneous recovery of fear. In Experiment 2 we first validated the IED procedure by showing that immediate extinction resulted in impaired extinction recall later during testing, compared to delayed extinction. We then found that cannabidiol given before immediate extinction reduced cued fear expression acutely without affecting baseline fear expression or extinction learning. Cannabidiol also resulted in enhanced extinction recall to rescue the IED, which required extinction and did not involve any effect on fear memory consolidation. Taken together, these results confirm previous findings demonstrating that cannabidiol reduces fear expression acutely and extends them by showing that cannabidiol protects against the return of fear after successful extinction and ameliorates stress-induced extinction resistance.

### Acute effects of cannabidiol on learned fear expression during delayed and immediate extinction

Our finding that cannabidiol reduced learned fear expression acutely during delayed or immediate extinction broadly agrees with previous results on cannabidiol regulation of contextual and cued fear expression. Studies have shown that systemic cannabidiol treatment reduces contextual fear expression [10-13]. We also showed previously that cannabidiol reduces the expression of baseline and cued fear during delayed extinction [6]. It is worth noting that we found different effects of cannabidiol on baseline and cued fear expression during delayed and immediate extinction in this study. Cannabidiol reduced baseline but not cued fear expression during delayed extinction, whereas cannabidiol had no effect on baseline fear but reduced cued fear expression during immediate extinction. Cannabidiol regulation of learned fear expression has been shown to depend on the strength of fear conditioning, such that it reduced contextual fear after stronger conditioning but had no effect with weaker conditioning [12]. In Experiment 1 we used weaker conditioning parameters in comparison to our previous study [6], whereas we used stronger conditioning parameters in Experiment 2 to induce the IED [23]. Compared to weaker conditioning, stronger conditioning resulted in no increase in freezing during fear learning and only marginally increased baseline freezing during extinction and its recall; however, tone-induced freezing was increased during extinction and extinction recall, providing evidence of stronger fear conditioning. Our finding that cannabidiol reduced cued fear expression after stronger but not weaker conditioning is therefore consistent with these previous results [12], although this does not explain cannabidiol reduction of baseline fear expression after weaker but not stronger conditioning. An alternative interpretation of this result is that cannabidiol reduced context generalization after weaker but not stronger conditioning, which is similar to the reported effect of cannabidiol when given after contextual fear conditioning [33].

### Effects of cannabidiol on delayed extinction and later spontaneous fear recovery

We found no acute effect of cannabidiol during delayed extinction or any lasting effect on extinction recall the next day. While this finding is congruent with the results of our previous study [6], others have shown that cannabidiol enhances contextual and cued fear extinction. Local cannabidiol infusion into the lateral ventricle or infralimbic cortex (IL) before repeated sessions of contextual fear extinction resulted in enhanced extinction recall [14-15]. The effects of cannabidiol on contextual fear extinction have also been shown to depend on the strength of fear conditioning. Systemic cannabidiol treatment before extinction learning enhanced the extinction of stronger contextual fear but impaired the extinction of weaker contextual fear [12]. Another study found that cannabidiol given before cued fear extinction had no effect during extinction learning or recall testing, as we found here and previously. However, cannabidiol given after extinction learning enhanced cued extinction recall, suggesting that it facilitated extinction memory consolidation [16].

The lack of any enduring effect of cannabidiol on extinction recall in the present study may have involved a floor effect, given the low freezing levels observed during extinction recall testing. In support of this idea, we found that cannabidiol prevented the spontaneous recovery of cued fear 21 days after extinction, despite its lack of effect on extinction recall. This provides evidence that cannabidiol enhanced extinction encoding and, to our knowledge, this is the first study to show that cannabidiol protects against the return of fear after extinction. Das et al. (2013) [16] did find that cannabidiol given before or after cued fear extinction learning tended to reduce later fear reinstatement, which is the return of fear that occurs with non-reinforced presentation of the shock after extinction [17], but this effect did not reach significance. It should be noted that cannabidiol also impairs fear memory reconsolidation to result in the lasting reduction of learned fear over time [34-35]. However, it is unlikely that our results can be explained by cannabidiol disruption of reconsolidation instead of facilitated extinction since we found low fear during extinction recall testing, which indicates successful fear extinction.

### Validation of the immediate extinction deficit procedure

Our finding that immediate extinction resulted in impaired extinction recall, compared to delayed extinction, is in general agreement with previous studies using the IED procedure [23, 25, 30, 36]. We found low baseline fear expression before extinction, with no differences between delayed and immediate extinction. This result has been reported in some [30, 36-37] but not other [30] previous studies. We also found no differences in cued fear throughout delayed and immediate extinction. Again, this finding agrees with some previous studies [25, 30], although others have shown increased [36] or decreased [37-38] cued fear expression during immediate extinction, compared to delayed extinction. These inconsistencies may involve methodological differences (e.g. fear conditioning parameters, interval between conditioning and extinction, rat strain, etc.) between studies. We did find higher baseline fear expression during extinction recall testing with immediate extinction, which reached significance in comparison to delayed no extinction. It is possible that the stress caused by recent conditioning in combination with tone-induced fear during immediate extinction induced an interoceptive state that became associated with the extinction context, resulting in a form of contextual fear conditioning [17]. We also found higher cued fear expression later on during extinction recall testing with immediate extinction, compared to delayed extinction, while there were no differences between immediate and no extinction. This adds to a growing number of studies that have demonstrated the IED [22].

### Effects of cannabidiol on the immediate extinction deficit

We found no acute effects of cannabidiol on extinction learning during immediate extinction. However, cannabidiol resulted in reduced baseline and cued fear expression during extinction recall testing, compared to vehicle, indicating that cannabidiol enhanced extinction recall to rescue the IED. This provides further evidence of cannabidiol enhancement of fear extinction by demonstrating its remediation of stress-induced extinction resistance, which is another novel finding of this study. Previous studies have shown that the IED results from high arousal or stress levels caused by recent fear conditioning. The IED fails to occur after weaker conditioning or with antagonism of receptor signalling mediated by various stress mediators [23-28]. This raises the possibility that cannabidiol ameliorated the IED at least in part by dampening the stress state associated with recent fear conditioning, although this remains to be determined in future studies.

The lack of effect of cannabidiol with no extinction indicates that cannabidiol required extinction to mediate its effect on immediate extinction, instead of directly having an enduring effect on fear expression during extinction recall. This is similar to a previous study showing that cannabidiol facilitation of the extinction of stronger contextual fear required extinction since it had no effect without extinction [12]. Our results also rule out an effect of cannabidiol on fear memory consolidation as a possible explanation for its effect on immediate extinction. Cannabidiol given systemically or infused into dorsal hippocampus reduces the consolidation of contextual fear [33, 39]. Taken together with our results, this raises the possibility that cannabidiol regulation of fear memory consolidation is specific to contextual fear. We also found evidence for extinction savings with immediate extinction since extinction recall was enhanced later on during testing using more tones, compared to no extinction, in vehicle-treated controls. This confirms previous results and suggests that immediate extinction impairs, rather than abolishes, fear extinction encoding [30].

### Conclusion

In this study we found that cannabidiol prevented the spontaneous recovery of fear after delayed extinction and ameliorated the extinction impairment that occurs with immediate extinction. Further research is needed to determine the pharmacological and neural mechanisms underlying cannabidiol regulation of the return of fear over time and stress-induced extinction resistance. Reduced contextual fear expression by cannabidiol depends on serotonin 5-HT1A receptors, whereas cannabidiol facilitation of contextual fear extinction is dependent on cannabinoid receptor type 1 (CB1R) signalling [6, 9]. Cannabidiol has low affinity for CB1Rs, suggesting that the CB1R dependency of its effect is mediated indirectly by modulating endocannabinoid transmission. This is supported by evidence indicating that cannabidiol increases the levels of brain endocannabinoids, which act as endogenous CB1R agonists [40]. Endocannabinoid-mediated CB1R transmission might therefore be involved in cannabidiol regulation of delayed extinction to prevent spontaneous fear recovery and of immediate extinction to rescue the IED. Cannabidiol may also regulate immediate extinction indirectly by reducing the stress arising from recent fear conditioning, which is mediated by adrenergic and corticotropin releasing factor (CRF) signalling in amygdala [25-28]. Cannabidiol blocks stress-induced increases in amygdala CRF expression [41], providing support for this idea. While evidence for the direct regulation of amygdala adrenergic transmission by cannabidiol is limited, this could also occur indirectly via elevated endocannabinoid levels. Previous studies have shown that endocannabinoid transmission regulates adrenergic signalling in amygdala [42-43] and other areas that comprise the fear extinction circuit, such as IL [44-45]. Regardless of the pharmacological and neural mechanisms involved, the combination of reduced fear expression and enhanced fear extinction makes cannabidiol an attractive candidate to investigate further as a potential adjunct to strengthen exposure-based therapy. This may help to achieve a lasting reduction in symptom relapse for the treatment of anxiety-related disorders.

## Materials and Methods

### Animals

Adult male Lister hooded rats (Charles River or Envigo, UK) weighing 200-480 g at the beginning of the experiments were used in this study. Rats were group housed (3-4/cage) in individually ventilated cages on a 12 h light/dark cycle (lights on at 8:00) with free access to food and water. Rats were humanely culled with a rising concentration of CO_2_ at the end of the experiments. All experimental procedures were conducted with ethical approval from the University of Nottingham Animal Welfare and Ethical Review Body and in accordance with the Animals (Scientific Procedures) Act 1986, UK (Home Office Project licence numbers 30/3230 and P6DA59444).

### Drug administration

Cannabidiol (THC Pharm, Germany) was suspended in vehicle (2% Tween 80 and sterile saline) on the day of use. In Experiment 1, cannabidiol (10 or 20 mg/kg) or vehicle was injected (i.p., 1 mL/kg) at a dose range used in previous studies that investigated its effects on fear extinction [6, 12]. In Experiment 2, one dose of cannabidiol (10 mg/kg) or vehicle was injected as above based on the results of Experiment 1.

### Behavioural testing

The behavioural testing procedures used were adapted from our [6, 29] and other [23, 30] previous studies on delayed and immediate extinction. Four behavioural testing chambers were used and the apparatus has been described elsewhere [31]. Tone and footshock presentations were controlled by a PC running MED-PC V software (Med Associates, US) and behaviour was recorded for later data analysis (see below). All behavioural testing was conducted during the rats’ light cycle.

In Experiment 1 we examined the effects of cannabidiol given before delayed extinction on extinction recall and the later spontaneous recovery of fear (Fig 1A). Rats were randomly allocated to receive one of the cannabidiol doses or vehicle. On Day 0, rats were habituated to two distinct contexts (A and B; 10 min each). On Day 1, rats underwent auditory fear conditioning in context A, consisting of five tones presented alone (30 s, 4 kHz, 80 dB, 2 min inter-trial interval (ITI)) followed by five pairings of the tone with footshock (0.5 s, 0.4 mA, ending at tone offset). On Day 2, rats were injected with drug or vehicle and 30 min later underwent extinction in context B, which consisted of 30 tones presented alone (30 s ITI). On Day 3, rats underwent extinction recall testing drug-free in context B, consisting of three tones presented alone (30 s ITI). On Day 23, rats underwent spontaneous fear recovery testing drug-free in context B, which also consisted of three tones presented alone (30 s ITI).

In Experiment 2 we first validated the IED procedure by comparing the effects of immediate and delayed extinction on later extinction recall (Fig 2A). Rats were randomly allocated to the four following groups: immediate extinction, delayed extinction, immediate no extinction, and delayed no extinction. All rats underwent auditory fear conditioning, consisting of five tones (as above) paired with footshock (1 s, 0.5 mA, ending at tone offset) in context A; stronger footshocks were used since weaker conditioning parameters fail to induce the IED [23]. Rats were returned to their home cage after fear conditioning. Immediate and delayed extinction occurred 30 min and 24 h after conditioning, respectively, and consisted of 45 tones presented alone (30 s ITI) in context B. Immediate and delayed no extinction also occurred 30 min and 24 h after conditioning, respectively, but the rats were placed in context B for the same duration without any tone presentations. All rats underwent extinction recall testing 24 hr after immediate, delayed, or no extinction, which consisted of five tones presented alone (30 s ITI) in context B.

We then examined the effects of cannabidiol given before immediate extinction on subsequent extinction recall to determine if cannabidiol rescues the IED (Fig 3A). Rats were randomly allocated to the four following groups: cannabidiol extinction, vehicle extinction, cannabidiol no extinction, and vehicle no extinction. All rats underwent auditory fear conditioning in context A as above, followed immediately by drug or vehicle treatment, after which the rats were returned to their home cage. Immediate and no extinction occurred 30 min after conditioning in context B as above. All rats underwent extinction recall testing drug-free 24 hr after immediate or no extinction, which consisted of 10 tones presented alone (30 s ITI) in context B.

### Data analysis

Freezing (i.e. absence of movement except in relation to respiration) in response to tone presentations during fear conditioning, extinction, extinction recall testing (Experiments 1-2), and spontaneous fear recovery testing (Experiment 1) was quantified as the learned fear response. Freezing was scored automatically using VideoTrack software (ViewPoint, France) as we have described previously [32]. The cumulative duration of freezing during the tones was calculated and expressed as a percentage of the tone duration. Baseline fear at the start of extinction, extinction recall testing (Experiments 1-2), and spontaneous fear recovery testing (Experiment 1) was inferred from freezing during the 2 min period before tone presentations and quantified as above.

In Experiment 1, the mean percentage of freezing in response to three consecutive tones during extinction was calculated for the statistical analysis. Similarly, the mean percentage of freezing in response to the three tones during extinction recall and spontaneous fear recovery testing was calculated for the statistical analysis. Differences in freezing in response to tone-shock pairings during fear conditioning were analyzed using two-way analysis of variance (ANOVA), with dose and trial as between- and within-subject factors, respectively. Differences in baseline freezing before extinction were analyzed using one-way ANOVA, with dose as the between-subjects factor. Differences in tone-induced freezing at the start of extinction were analyzed in the same way to examine cued fear expression. Differences in tone-induced freezing during extinction were analyzed using two-way ANOVA, with dose and trial block as between- and within-subjects factors, respectively. Differences in baseline freezing during extinction recall and spontaneous fear recovery testing were analyzed using two-way ANOVA, with dose and time as between- and within-subjects factors, respectively. Differences in tone-induced freezing during extinction recall and spontaneous fear recovery testing were analyzed in the same way.

In Experiment 2, the mean percentage of freezing in response to five consecutive tones during extinction was calculated for the statistical analysis. The mean percentage of freezing in response to the five (validation) or 10 (cannabidiol) tones during extinction recall was also calculated for the statistical analysis. Differences in freezing in response to tone-shock pairings during fear conditioning were analyzed using two-way ANOVA as above. For validation of the IED procedure, baseline freezing data before extinction from the two immediate (extinction and no extinction) and two delayed (extinction and no extinction) groups were combined. For determining the effects of cannabidiol on the IED, baseline freezing data before extinction from the two cannabidiol (extinction and no extinction) and two vehicle (extinction and no extinction) groups were combined. Differences in baseline freezing before extinction were analyzed using two-tailed unpaired t-tests. Differences in tone-induced freezing at the start of extinction were analyzed in the same way to examine cued fear expression. Differences in tone-induced freezing throughout extinction were analyzed using two-way ANOVA as above. Differences in baseline freezing during extinction recall testing were analyzed using one-way ANOVA, with group as the between-subjects factor. Differences in tone-induced freezing during extinction recall testing were analyzed in two ways. Differences in mean freezing across the five (validation) or 10 (cannabidiol) tones were analyzed using one-way ANOVA, with group as the between-subjects factor. Differences in freezing in response to each tone were also analyzed using two-way ANOVA, with group and trial as between- and within-subject factors, respectively.

All data are presented as the mean + standard error of the mean. Post-hoc comparisons were conducted using the Newman-Keuls or Sidak’s tests where indicated. The level of significance for all comparisons was set at P < 0.05.

## Acknowledgements

Cannabidiol was generously supplied by Artelo Biosciences.

## Author contributions

EP, WW, and CS: Conceived the study, designed the experiments, and analyzed the data. EP and WW: Conducted the experiments. EP, WW, HC, and CS: Drafted the paper. All authors contributed to manuscript revision and approved the submitted version.

## Institutional review board statement

All experimental procedures were conducted with ethical approval from the University of Nottingham Animal Welfare and Ethical Review Body and in accordance with the Animals (Scientific Procedures) Act 1986, UK (Home Office Project licence numbers 30/3230 and P6DA59444).

## Data availability statement

The data presented are available on reasonable request from the corresponding author.

